# Dual Inhibition of Cathepsin L and 3CL-Pro by GC-376 Constrains SARS Cov2 Infection Including Omicron Variant

**DOI:** 10.1101/2022.02.09.479835

**Authors:** Prabhakaran Kumar, Kiira M Ratia, Justin M Richner, Gregory R J Thatcher, Rashmi Kadam, Sandra P Smieszek, Bartlomiej P Przychodzen, Vuk Koprivica, Gunther Birznieks, Mihael H Polymeropoulos, Bellur S Prabhakar

## Abstract

Recurrent waves of SARS CoV2 infections remain a major global health concern. Emergence of highly infectious variants with reduced sensitivity to neutralization by vaccines and monoclonal antibodies (mAb) necessitates a deeper understanding of factors involved in SARS CoV2 infections and identification of drug candidates to halt infection. Here, we determined the primacy of endosomal protease cathepsin-L in mediating SARS CoV2 entry and screened a library of well-annotated bioactive compounds for potent cathepsin-L inhibitory activity. Whilst the potent cathepsin-L inhibitors were capable of inhibiting SARS CoV2 entry and cytopathic effect (CPE) in less susceptible cell lines such as human ACE2 expressing 293T cells, these drugs failed to inhibit SARS CoV2 in highly susceptible cell lines such as human TMPRSS2 or human-ACE2-TMPRSS2 overexpressing Vero E6 cells. Only drugs with dual inhibitory effect on both host cathepsin-L and virus 3CL-Protease enzymes such as Z-FA-FMK and GC-376 were capable of inhibiting prototypic (USA-WA1/2020, Lineage A) SARS CoV2 induced CPE in highly susceptible cell lines. Moreover, these drugs inhibited delta (Lineage-B.1.617.2) and omicron (Lineage-B.1.1.529) infection with equal potency showing that the newer mutations harbored in these variants did not affect the mechanism of action of these drugs such as cathepsin-L or 3CL-Pro inhibition. Moreover, our early evidence that 3CL-Pro inhibition can effectively inhibit omicron-induced CPE in highly susceptible cell lines suggests that the recently FDA-approved oral drug, a 3CL-Pro inhibitor which is a combination of nirmatrelvir/ritonavir (Paxlovid) could be effective against omicron variant which shows reduced sensitivity to vaccines and mAb.

**Importance:** We report that cathepsin-L and 3CL-Pro as major targets for designing antivirals against SARS CoV2. Dual inhibition of cathepsin-L and 3CL-Pro by GC-376 renders it effective in inhibiting SARS CoV2-induced cytopathic effect in highly susceptible cell lines. Moreover, this candidate drug is equally effective against prototypic SARS CoV2 lineage A and emerging variants such as delta and omicron which show reduced sensitivity to vaccines and monoclonal antibodies. Given the recent wave of SARS CoV2 omicron variant infection around the world, and 3CL-Pro inhibitor nirmatrelvir is one of the components of the FDA-approved Paxlovid, our findings are timely, important and should be of broad interest.

## Introduction

COVID19, originating in the Hubei province of China, is one of the most devastating pandemics the world has ever witnessed (1). The repeated waves of COVID19 have caused more than 370 million infections and accounted for 5.65 million deaths so far. While vaccination is proving to be the most effective weapon to fight this pandemic, scaling up the production and distribution to meet the global need is extremely challenging. Poor vaccination rates in third world countries can result in highly infectious forms of the variants such as delta and omicron variants which might necessitate administration of booster doses of vaccine (2). On the other hand, vaccine hesitancy could facilitate the spread of these newer variants despite the significant progress in vaccinations.

A major concern associated with COVID19 is the emergence of mutant strains with higher transmissibility and improved immune evasion mechanisms. For instance, the delta variant (B.1.617.2) was found to have ~2-fold higher transmissibility than the prototypic lineage-A strain. The delta variant was shown to encompass 9 mutations within the spike protein which is targeted by many of the available vaccine candidates (3) and was found to be 2.9 times less susceptible to neutralization by convalescent serum from recovered COVID19 patients (infected with lineage A) and sera from vaccinated individuals compared to the lineage A USA/WA1/2020 strain (3). Similarly, increased immune evasion and reduced sensitivity to neutralization by the sera from vaccinated individuals and monoclonal antibodies (mAb) have also been reported with the omicron variant which has 30 mutations in the spike region (4, 5). The reduced neutralizing activity can be correlated with the mutations harbored within the spike protein. All combinations of the FDA-approved mAbs such as tixagevimab plus cilgavimab, imdevimab plus casirivimab, and etesevimab plus bamlanivimab, which neutralized the early strain and the alpha and delta variants were found to be less effective in neutralizing the omicron variant in at least *in vitro* neutralization assays (6).

SARS-nCoV-2 is a member of the Coronaviridae family and is an enveloped single-stranded RNA beta coronavirus. The RNA genome of SARS CoV2 encodes 4 structural proteins namely, spike (S), membrane (M), envelope (E), and nucleocapsid (N) proteins, and 16 nonstructural proteins (NSPs). The proteolytic processing is carried out by two virus proteases namely, Papain-Like Protease (PL-Pro) and 3C-Like protease (3CL-Pro alias M-Pro) which process NSP1-4 and NSP5-16 respectively (7). RNA-dependent RNA polymerase (RdRp) of SARS Cov2 comprises of a catalytic subunit (nsp12) and two auxiliary subunits, (nsp8 and nsp7) which regulate the replication and the transcription of its genes (8). Conventional antiviral drugs target specific events of the virus life cycle such as viral attachment, fusion, replication, and protein synthesis. As of now, FDA has granted authorization for three antiviral drugs namely remdesivir, molnupiravir, and nirmatrelvir/ritonavir. Among these, remdesivir and molnupiravir are inhibitors of RNA-dependent RNA polymerase (RdRp), and nirmatrelvir targets SARS CoV2 main protease (3CL-Protease) (9) (10). Apart from the mutations located within the spike protein, Omicron variant has the second highest number of non-synonymous mutations in the RdRp. Additionally, mutations were also observed in the 3CL-protease encoding regions (6, 11) which raise concerns about the clinical efficacy of these antiviral drugs.

Human ACE2 has been characterized as the most predominant receptor for SARS CoV2 (12, 13). The cell entry mechanisms of enveloped viruses such as coronaviruses require the attachment of the viral spike (S) structural glycoprotein to the host receptor and fusion with the host cell membrane which requires proteolysis and acidification in endosomes (14). Upon binding to ACE2 receptor, SARS CoV2 exploits host proteases such as transmembrane serine protease 2 (TMPRSS2) (13) and lysosomal cysteine proteases such as cathepsins for the cleavage of the spike protein into S1/S2 subunits followed by fusion of S2 subunit to the host cells (12, 15). Intriguingly, SARS CoV2 spike protein contains an additional furin cleavage site, which is absent in SARS CoV1(16) but present in MERS CoV (17), to exploit an additional host factor enhancing its infectivity. Contradicting reports suggest varying degrees of importance of these host proteolytic enzymes in regulating SARS CoV2 entry. Earlier, we have shown that small molecules that can prevent cathepsin L cleavage of viral glycoproteins derived from SARS-CoV and Ebola, Hendra, and Nipah viruses could effectively inhibit their entry into the host cells (18).

In this present study, we investigated the relative importance of TMPRSS2 and cathepsin L in mediating SARS CoV2 entry and determined that cathepsin L is more critical in mediating SARS CoV2 entry. Further, we screened a commercially available bioactive compound library for cathepsin L inhibitory activity and evaluated the ability of these inhibitors to prevent the entry of pseudotyped SARS CoV2 particles as well as to inhibit infectious SARS CoV2 induced cytopathic effect (CPE) in various cell lines. Our screening identified two known protease inhibitors with pronounced cathepsin L inhibitor activity, namely Z-FA-FMK and GC-376, which effectively inhibited virus entry and CPE at sub-micro-molar doses in 293T cells overexpressing human ACE2. Both these compounds inhibited delta and omicron variants-induced CPE with the same efficacy as they inhibited prototypic USA-WA1/2020 strain. More importantly, GC-376 which is a potent inhibitor of 3CL-Pro inhibited SARS CoV2 induced CPE in highly susceptible Vero cells overexpressing human ACE2 and TMPRSS2 indicating its higher potency.

## Materials and Methods

### Cathepsin-L primary assay

The cathepsin-L primary assay measures protease activity with a peptide substrate derived from the SARS-CoV-2 Spike protein. The FRET-based peptide, (TAMRA)-IIAYT/MSLGAK-(FAM), was synthesized by Anaspec. All cathepsin-L assays were performed at room temperature in black, flat-bottom 384-well plates containing a final reaction volume of 50 μL. Reactions were assembled as follows: 40 uL of 6.25 μg/mL human CatL (Novoprotein) in Buffer A (50 mM ammonium acetate, pH 5.5, 0.1 mg/mL BSA, 4 mM EDTA, 2.5 mM TCEP, 0.1% (v/v) Pluronic F-68) was dispensed into wells containing 0.1 μL of test compound in DMSO or appropriate controls. The enzyme was incubated with the compounds for 10 min before substrate addition. Reactions were initiated with 10 μL of 10 μM FRET peptide in Buffer A. Plates were shaken vigorously for 30 s, and fluorescence generated by peptide cleavage was monitored continuously for 15 min on a Tecan Infinite M200 Pro plate reader (λ_excitation_=485 nm; λ_emission_=535 nm). Slopes from the linear portions of each progress curve were recorded and normalized to plate-based controls. Positive control wells, representing 100% inhibition, excluded cathepsin-L enzyme; negative control wells, representing 0% inhibition, included vehicle.

### Cathepsin L high-throughput screening (HTS)

HTS for inhibitors of CatL was performed using the primary assay, described above, against the TargetMol Bioactives Library. The library consists of 5,370 compounds with known biological activities and includes clinical, pre-clinical, and approved drugs. Test compounds were screened at 20 uM final concentration and were delivered via 100 nL pin tool (V&P Scientific) alongside DMSO controls. Each 384-well plate contained 32 positive control wells, 32 negative control wells, and 320 test compound wells. The average plate-based Z’ value for this assay was 0.74. Compounds producing >50% inhibition of CatL activity were selected for follow-up analysis.

### CRISPR CAS9 deletion of human CTSL and TMPRSS2

Guide RNAs targeting CTSL gene (5’ CAGTATGTTCAGGATAATGG 3’ and 5’ AGATGTTCCGGAAAACTGGG 3’) and 5’ ACTGGAACGAGAACTACGGG 3’ and 5’ GGATGAAGTTTGGTCCGTAG 3’ were picked from Broad institute website (https://portals.broadinstitute.org/gpp/public/analysis-tools/sgrna-design). Synthesized guide RNA, TracrRNA, and Cas9 enzyme were purchased from IDT, Inc. Briefly, 1 x 10^6 293T ACE2 cells were nucleofected with sgRNA:TracrRNA: Cas9 complex using Amaxa Cell Line Nucleofector Kit V and cultured for 72h. Single-cell clones were sorted and screened for deletion of the targeted region by PCR using the following primer sets: CTSL Forward 5’ AGCCAAGAAGTGATTGACATCTTT3’ and reverse 5’CTAATATATGAATTCCGAAAAAGGTA3’; TMPRSS2 Forward 5’GAATCACTTGAACCTGGGAGGCGGAG and reverse 5’GCTGGGCACAGCCCTAGCAGGACA3’. All experiments were approved and performed per the guidelines set forth by the Institutional Biosafety Committee (IBC) and the Environmental Health and Safety Office (EHSO) at the University of Illinois at Chicago.

### SARS CoV2 spike protein cleavage assay

Recombinant SARS-CoV-2 Spike protein with C-terminal 6-His tag was purchased from R & D Systems, Inc. 500ng of the spike protein was treated with 100ng recombinant human cathepsin-L (Sigma-Aldrich) in sodium acetate buffer (pH 5.5) at room temperature for 4h in the presence of 2 and 20 μM concentrations of indicated inhibitors. Cleavage of the spike protein was detected using anti-SARS-CoV Spike Protein (SDelta3) monoclonal antibody (7G12, Thermo Fisher Scientific, CA, USA)) raised against immunogen aa168-461 of the S1 domain.

### Pseudovirus entry inhibition assay

SARS CoV2 pseudotyped virus particles were produced by transfecting HEK293T cells with the following plasmids 1) lentiviral backbone plasmid-pHAGE-CMV-Luc2-IRES-ZsGreen that expresses luciferase and ZsGreen reporters.2) pHDM-Hgpm2-lentiviral helper plasmid expressing HIV Gag-Pol. 3) pHDM-tat1b: lentiviral helper plasmid expressing HIV Tat.4) pRC-CMV-Rev1b: lentiviral helper plasmid expressing HIV Rev, and pHDM expressing the SARS CoV2 Wuhan-Hu-1 Spike envelope Glycoprotein (Procured from BEI resources). Bald pseudovirus particles were produced as described above without spike glycoprotein. 25 x 10^3^ human ACE2 expressing 293T cells were seeded in 96 well plates and infected with bald pseudovirus without spike glycoprotein (negative control) and SARS CoV2 spike protein-expressing pseudovirus and treated with 1 μM concentration (50μl Infection medium 2% FBS + 150 μl pseudovirus) of the candidate inhibitory small molecules. 72 hours later, the cells were lysed and the luciferase activity was determined by luciferase assay. Briefly, the spent medium was removed and 60 μl of the cell lysis buffer reagent (Promega Inc.) was added to each well. Plates were incubated for 15 min with shaking. 40 μl of luciferin substrate was added and luminescence read with 1 min integration and delay time. The percentage inhibition was calculated as follows: 100 x [1 - (X - MIN)/(MAX - MIN)]. X= Test RLU; Min = RLU of Mock without envelope; Max = RLU of infected control.

### CPE assay using infectious SARS CoV2

25 x 10^3^ hACE2-293T (BEI resources) and human TMPRSS2 expressing Vero E6 cells (Ordered from Japanese Collection of Research Bioresources Cell Bank (JCRB) and distributed by Sekisui XenoTech, LLC, KS) or Vero E6 cells (ATCC), Vero E6 Cells Expressing TMPRSS2 and Human ACE2 (Vero E6-TMPRSS2-T2A-ACE2) (NR-54970, BEI) were seeded in each well of 96 well plates in 100 μl of 5% DMEM medium. SARS-CoV-2, Isolate USA-WA1/2020, Isolate hCoV-19/USA/PHC658/2021 (Lineage B.1.617.2; Delta Variant; NR-55611) and Isolate hCoV-19/USA/MD-HP20874/2021 (Lineage: B.1.1.529, Omicron Variant BEI Cat # NR-56461) obtained from BEI resources were used to infect the cells. 100 μl of the infection medium (2% DMEM) containing 100 x TCID50 virus was used to infect cells. The assay was performed in triplicates with uninfected/infected cells with or without inhibitors (200nM, 1μM, and 5 μM). CPE was quantified at 72 h post-infection using ATPLite luminescence assay. Briefly, 100 μl of spent medium was removed and 100 μl of the Cell Titer Glo assay reagent (Promega Inc.) was added to each well. Plates were incubated for 15 min with shaking. Luminescence was read with 1 min integration and delay time. The virus-induced cytotoxicity % was calculated as follows: 100 x [1 - (X - MIN)/(MAX - MIN)].X = Test RLU; Min = RLU of infected control wells; Max = Mean RLU of uninfected control wells.

### 3CL-Pro and PL-Pro activity assays

Inhibitory effect of selected compounds from 0.5 nM to 50 nM concentrations were determined using 3CL pro and PL-Pro assay kits (BPS Bioscience, CA, USA) as per manufacturer’s instructions. Briefly, indicated doses of the compounds were pre-incubated with 150ng of 3CL-Pro or 15ng of PL-Pro for 30 min and then with 50μM fluorogenic substrate either overnight at room temperature (for 3CL-Pro) or 1 hour at 37°C (for PL-Pro). The fluorescence at 460nM intensity was measured using a microplate reader.

### Statistical analysis

Statistical analyses were performed using Prism GraphPad (V9.0). Data were expressed as Mean ± SEM. Student’s t-test was used to compare two groups, whereas ANOVA with Tukey’s multiple comparisons was used to compare more than two groups. A p-value < 0.05 was considered significant.

## Results

### 1. Cathepsin L cleaves SARS CoV2 spike and is indispensable for SARS CoV2 entry in 293T ACE2 cells

We validated the use of hACE2 overexpressing 293T (hACE2-293T) cells for studying SARS CoV2 pseudovirus entry. We compared the specificity of SARS CoV2 spike-hACE2 mediated pseudovirus entry by transducing wild type 293T vs hACE2-293T cells with VSV glycoprotein and SARS CoV2 spike glycoprotein pseudotyped virus particles produced with luciferase reporter vector. As shown in Fig-1A-B, VSV-G pseudotyped virus particles efficiently entered into both WT 293T and 293TACE2 cells, whereas SARS CoV2 pseudotyped virus particles entered only in hACE2-293T cells but not WT 293T cells. We performed an *in vitro* S1/S2 cleavage assay in the presence of functional human CTSL enzyme and confirmed the ability of CTSL to cleave SARS CoV2 spike (Fig-1C).

**Figure-1:**
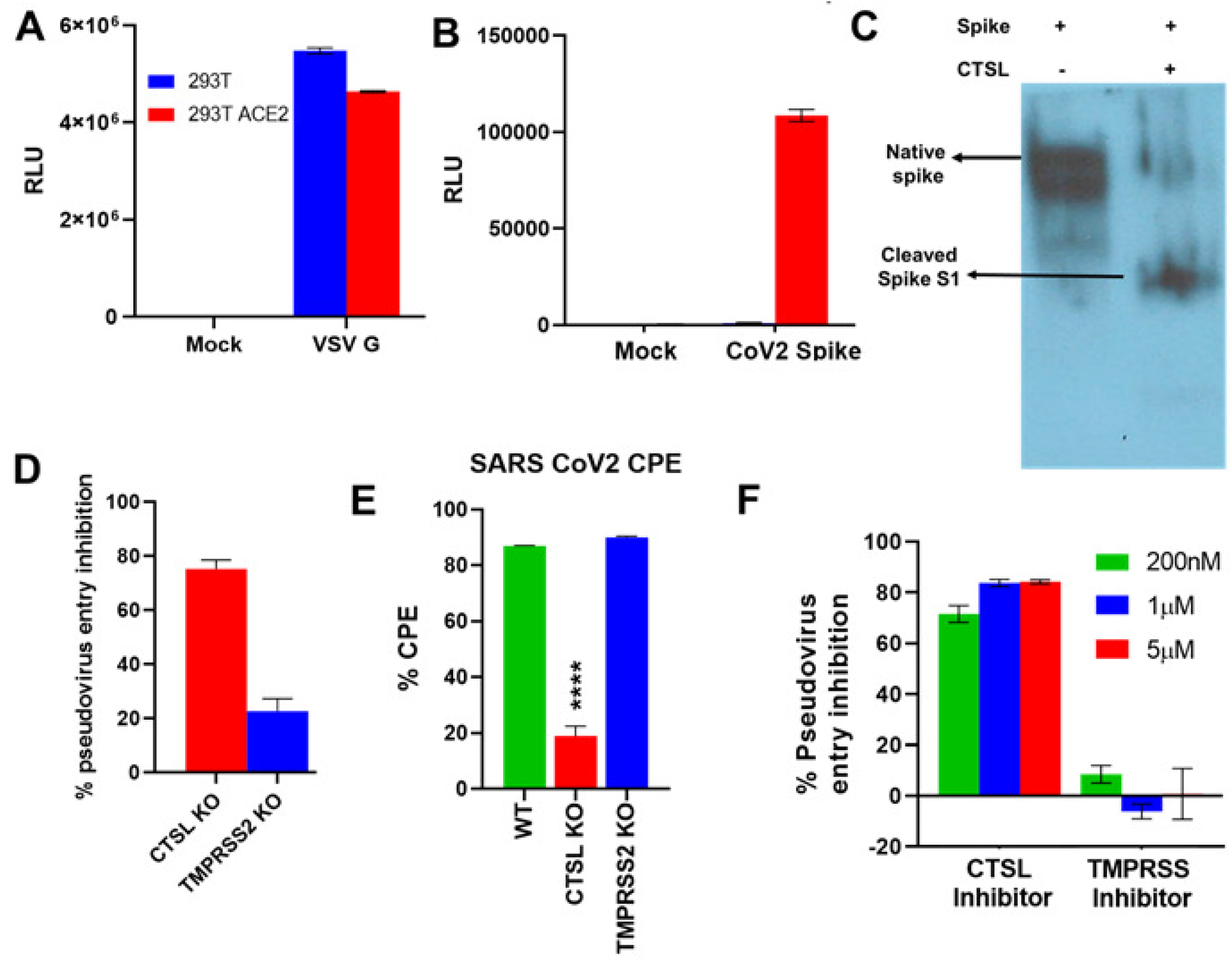
A-B) 293T WT and hACE2 overexpressing 293T (293T ACE2) cells were transduced with VSV-G glycoprotein (A) and SARS CoV2 spike protein (B) pseudotyped virus particles and bar graphs show RLU values for each cell type. C) Western blot shows cleavage of SARS CoV2 spike protein by CTSL enzyme. D) 293T ACE2, CTSL and TMPRSS2 deleted 293T ACE2 cells were transduced with SARS CoV2 spike protein pseudotyped virus particles and the bar graph shows relative pseudovirus entry inhibition compared to WT 293T ACE2 cells. E) Bar graph shows % inhibition of SARS CoV2 induced cytopathic effect (CPE) in 293T ACE2, CTSL and TMPRSS2 deleted 293T ACE2 cells 72h post-infection. (n=3, **** p<0.0001 Vs WT) F) Bar graph shows % inhibition of SARS CoV2 pseudotyped virus entry in 293T-ACE2 cells exhibited by the pharmacological inhibitors of CTSL and TMPRSS enzymes. (Assay was done in triplicates and two independent experiments. Values are expressed as Mean ± SEM).

To determine the relative importance of human CTSL and TMPRSS2 in mediating SARS Cov2 entry in 293T ACE2 cells, we knocked out the expression of these genes by using the CRISPR-Cas9 method, and the single-cell clones were expanded. The deletion of the targeted exon was confirmed by RT-PCR (Fig-S1). Next, we determined the effect of the deletion of the CTSL and TMPRSS2 genes on SARS CoV2 virus entry into 293T-ACE2 using the pseudovirus entry assay. Intriguingly, SARS CoV2 pseudotyped virus entry into CTSL deleted 293T ACE2 cells was inhibited by ~70-80% relative to WT-293T ACE2 cells. On the other hand, only ~20% of pseudovirus entry inhibition was noted in TMPRSS2 deleted 293T ACE2 cells suggesting a dominant role of CTSL in mediating SARS CoV2 entry in 293T ACE2 cells compared to TMPRSS2 (Fig-1D).

Next, we determined the susceptibility of WT, *CTSL^KO^*, and *TMPRSS2^Ko^* 293T-ACE2 cells to SARS CoV2 infection. We monitored virus-induced cytopathic effect (CPE) using a luminescence-based cell viability assay and found that CTSL deleted 293T-ACE2 cells were ~75% less susceptible to SARS CoV2 induced CPE compared to WT cells, whereas there was no significant reduction in the CPE in TMPRSS2 KO cells (Fig-1E). Additionally, we determined the relative effect of these proteases using pharmacological inhibitors. As shown in Fig-1F, cathepsin–L inhibitor effectively inhibited pseudovirus entry (>50%) at all doses tested ranging from 200nM to 5μM, whereas TMPRSS inhibitor did not show any inhibitory activity (Fig-1F).

### 2. FRET-based HTS Screening of cathepsin-L inhibitors

To identify inhibitors of cathepsin L-mediated cleavage of SARS-CoV2 Spike protein, we designed a FRET-based polypeptide containing the CTSL P5-P5’ recognition sequence within the spike protein: (TAMRA)-**IIAYT**/**MSLGA**K-(FAM). Peptide hydrolysis by purified CTSL enzyme could be followed continuously by measuring the appearance of FAM fluorescence over time, and hydrolysis could be inhibited by the addition of previously characterized CTSL inhibitors, such as triapine (data not shown). This assay was used to screen 5,370 compounds in the TargetMol Bioactives library, a collection of heavily annotated small molecules, including clinically-tested and FDA-approved compounds. The screen, performed using 20 uM test compound, returned an average Z’-value of 0.74 and identified 112 compounds that reduced the slope of the CTSL-catalyzed reaction by 50% (Figure 2A). Compounds that interfered with the assay signal or scored frequently in screens of unrelated targets were removed from further consideration. The remaining 95 compounds were triaged using a combination of factors, favoring: higher % CTSL inhibition; efficacy as anti-infective; activity as protease inhibitor; chemical diversity; and including representative anti-inflammatory agents.

**Figure-2:**
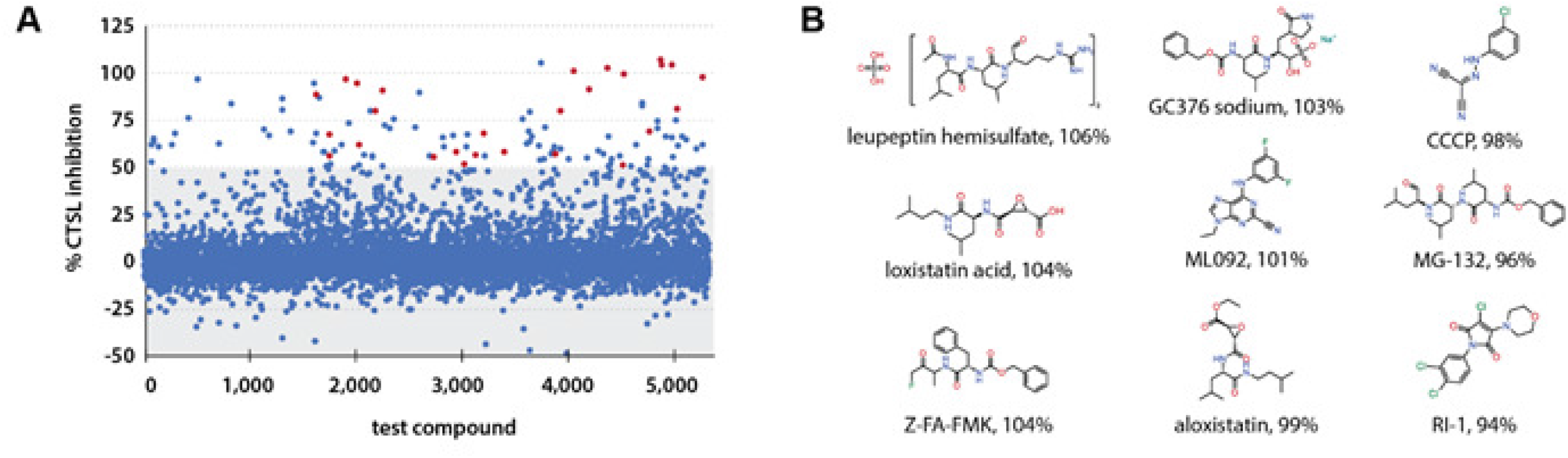
CTSL HTS of bioactive small molecules. A) A scatter plot of the inhibitory activity of each of the 5370 assayed compounds on CTSL activity with a FRET-based peptide representing a portion of the SARS CoV2 Spike protein. The grey section underlays the compounds that did not reach the cut-off criteria of 50% CTSL inhibition. The compounds that progressed to validation in the pseudovirus entry assay are represented by red circles. B) The structures of the top nine hits identified in the screen and prioritized for follow-up studies.

Bioactive libraries include many PAINS compounds (19) and although some were triaged because of assay interference, hits such as the quinone visomitin, a redox-active PAINS compound, were also actively triaged. Of the 27 hits selected, several could be considered as PAINS compounds because they contain an electrophilic or redox-active functional group that interacts with the active site cysteine of cysteine proteases and other enzymes: approximately 20 of the selected 27 hits have the potential for such interactions with the active site cysteine of CTSL. For example, MG-132 is an aldehyde 20S proteasome inhibitor that also inhibits calpains; boceprevir and telaprevir are ketoamide viral protease inhibitors; and GC-376 is a masked aldehyde electrophile, requiring bio-activation to inhibit the 3CLpro main cysteine protease of SARS-CoV-2(20).

### 3. Validation of HTS selected hits in pseudovirus entry inhibition and infectious SARS CoV2 CPE inhibition screening in 293T ACE2 cells

Next, 27 selected hits from our HTS CTSL cleavage assay were subjected to pseudovirus entry inhibition assay using 293T-hACE2 cells. Among them, leupeptin hemisulfate, loxistatin acid, Z-FA-FMK, aloxistatin, ML-092, CCCP, MG-132, balicatib, FCCP, GC-376 sodium, AVN944, triapine, GlyH-101, and ML-364 exhibited >50% pseudovirus entry inhibition in at least one of the tested doses i.e. 200nM, 1μM and 5μM (Fig-3A). We noted >50% cytotoxicity for compounds MG-132, AVN944, and triapine in 293T ACE2 cells (Fig-3B), and therefore, the observed reduction in RLUs in pseudovirus entry could be due to the cytotoxicity as well.

**Figure-3:**
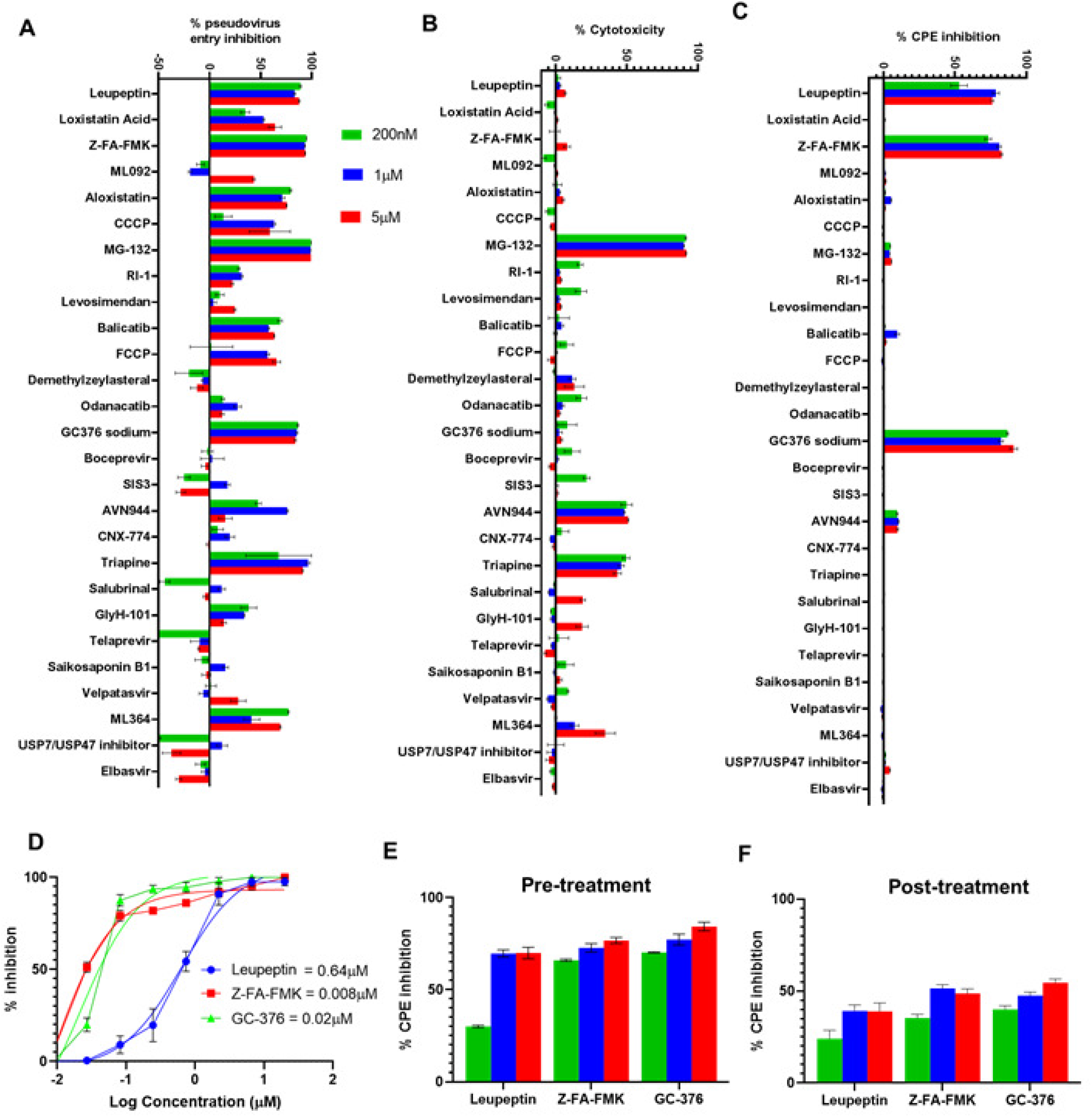
A) Dose-response effect of selected hits from the bioactive library on SARS CoV2 pseudotyped virus in 293T ACE2 cells. B) Drug-induced cytotoxicity of the selected hits from the bioactive library in 293T ACE2 cells. C) Dose-response effect of selected hits from the bioactive library on SARS CoV2 induced cytopathic effect in 293T ACE2 cells. D) CPE inhibition IC50 determination for leupeptin, Z-FA-FMK, and GC-376 sodium in 293T ACE2 cells. E-F) Effect of the pre (E) and post-treatment (F) on SARS CoV2 induced CPE in 293T ACE2 cells. (Assay was done in triplicates and three independent experiments. Values are expressed as Mean ± SEM).

Next, we tested the ability of these drugs to inhibit SARS CoV2 infection and the associated cytopathic effect in 293T-hACE2 cells. Intriguingly, only three drugs namely, leupeptin hemisulfate, Z-FA-FMK, and GC-376 sodium effectively inhibited SARS CoV2 induced CPE in 293T ACE2 cells, whereas none of the other drugs which inhibited pseudovirus entry in 293T ACE2 cells were capable of inhibiting cytopathic effect in 293T ACE2 cells (Fig-3C). IC50 determination experiments showed an IC50 value of 0.64μM, 0.008 μM, and 0.02 μM for leupeptin hemisulfate, Z-FA-FMK, and GC-376 respectively for inhibition of CPE in 293T ACE2 cells (Fig-3D).

Next, we tested the effect of these drugs on CPE when added 1h post-infection (after virus absorption). All three compounds had 25-20% reduction in their inhibitory effect when added post-infection (Fig-3E-F), but were still capable of producing more than 50% inhibitory effect on SARS CoV2 induced CPE.

### 4. GC-376 inhibits USA-WA1/2020, Delta and Omicron Variant SARS CoV2 induced CPE in highly susceptible cell lines with similar potency

While 293T ACE2 cells are permissive to SARS CoV2 entry, the CPE was induced only at lower dilution (10^-2^), but not at higher dilutions of the virus. On the other hand, Vero E6 cells are routinely used in SARS CoV2 CPE assays. However, drugs such as chloroquine/hydroxychloroquine that showed inhibitory effect of SARS CoV2 infection in Vero E6 cells (21) failed to exert an inhibitory effect on other highly susceptible cell lines such as Vero E6 cells overexpressing human TMPRSS2 (22) which is consistent with its failure in clinical trials (23). Therefore, a more rigorous selection of appropriate cell lines is key to identifying compounds with a higher probability of success in clinical trials. We compared SARS CoV2 induced CPE in 293T-ACE2, Vero E6 cells overexpressing human TMPRSS2 (Vero TMPRSS2) and Vero E6-overexpressing human ACE2 and TMPRSS2 (Vero ACE2-TMPRSS2). As shown in Figure-4A, 293T-ACE2 cells were the least susceptible to SARS CoV2 induced CPE with a TCID50 value of 0.007. Vero TMPRSS2 showed a TCID50 value of 2.87e-006 and the Vero ACE2-TMPRSS2 cells were the most susceptible to infection with a TCID50 value of 7.37e-007. Thus, overexpression of ACE2 and TMPRSS2 dramatically increased SARS CoV2 infectivity and CPE in Vero E6 cells. Further, we reaffirmed the CPE assay results with standard plaque assay where we found an increase in the number and size of plaques in cells expressing either TMPRSS2 or ACE2-TMPRSS2 compared to Vero E6 cells which were less susceptible to SARS CoV2 induced CPE (Fig-4B).

**Figure-4:**
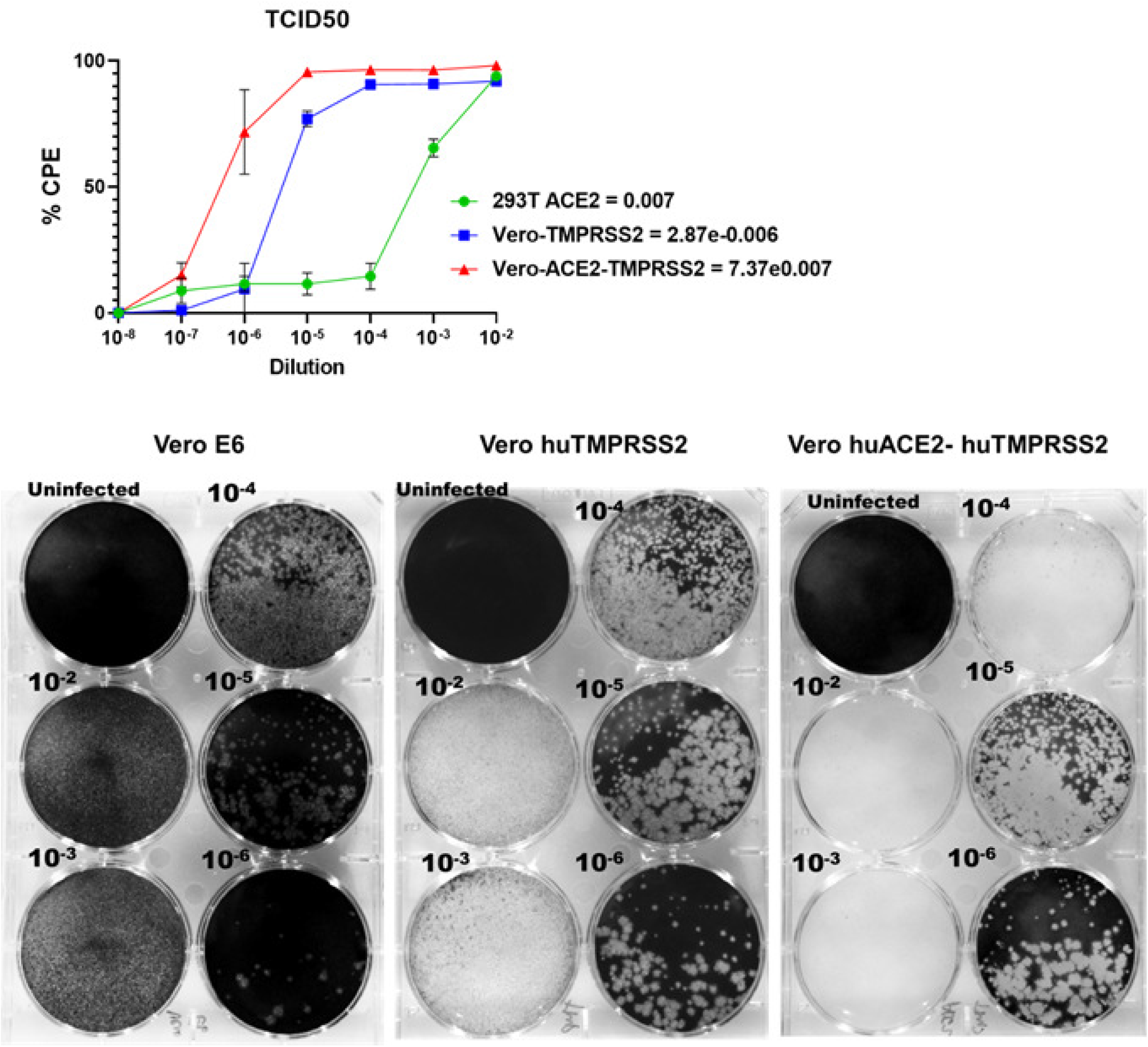
A) Sensitivity of different cell lines to SARS CoV2 induced CPE as determined by CPE assays at different dilutions. B) Vero E6, human TMPRSS2, and huACE2-TMPRSS2 overexpressing Vero E6 cells were infected with indicated dilutions (10^-2^-10^-6^) of SARS CoV2 isolate USA-WA1/2020, and plaques were developed with crystal violet staining 48h post-infection.

Next, we examined the effect of the candidate drugs to inhibit SARS CoV2 infection in Vero TMPRSS2 and Vero ACE2 TMPRSS2 cells. Compounds Z-FA-FMK and GC-376 inhibited SARS CoV2 CPE in Vero TMPRSS2 cells under both pre and post-treatment conditions with similar efficacy. GC-376 had a better inhibitory effect than Z-FA-FMK at 5μM dose (46.17±1.19 Vs 77.15±4.90). However, leupeptin hemisulfate failed to show significant inhibition of SARS CoV2 induced CPE in these cells under both pre and post-treatment conditions (Fig-5A-B). Both leupeptin hemisulfate and Z-FA-FMK were not effective in inhibiting SARS CoV2 induced CPE in the Vero-ACE2-TMPRSS2 cell line (Fig-5C-D). Only GC-376 inhibited SARS CoV2 CPE in Vero-ACE2-TMPRSS2 cells at the highest 5μM concentration (Fig-5C-D).

**Figure-5:**
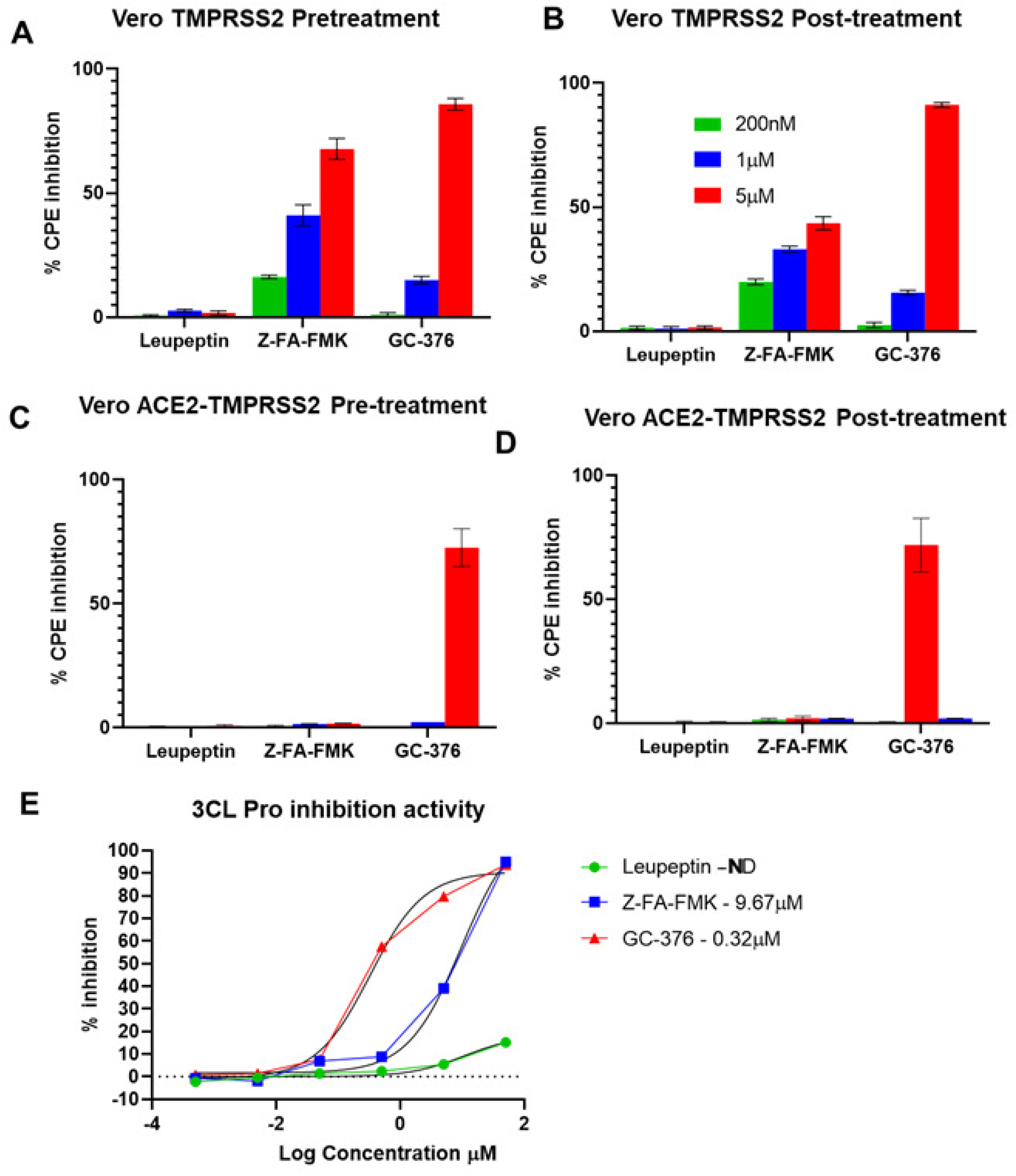
Effect of pre-treatment of leupeptin hemisulfate, Z-FA-FMK, and GC-376 on SARS CoV2 induced CPE in (A) Vero-huTMPRSS2 (B) Vero-huACE2-huTMPRSS2 cells. C-D) Effect of post-treatment of leupeptin hemisulfate, Z-FA-FMK, and GC-376 on SARS CoV2 induced CPE in (C) Vero-huTMPRSS2 and (D) Vero-huACE-huTMPRSS2 cells. E) IC50 determination for 3CL-Pro inhibitory activity of the compounds tested by *in vitro* fluorogenic substrate cleavage assay. (Assay was done in triplicates and three independent experiments. Mean ± SEM values from one independent experiment are shown).

While these drugs can effectively inhibit host proteases for preventing SARS CoV2 entry their effect on virus proteases such as 3CL pro and PLpro which are essential for virus replication couldn’t be ruled out. Therefore, we tested the efficacy of these compounds to inhibit 3CL-Pro and PL-Pro *in vitro* using fluorogenic substrate cleavage assay and found that leupeptin had no significant inhibitory effect on either PL-pro or 3CL-Pro. GC-376 and Z-FA-FMK had IC50 values of 0.32μM and 9.67μM respectively against 3CL-Pro (Fig-5E) they were ineffective in inhibiting PL-Pro (data not shown). Thus, it is clear that the differences in the antiviral activity of GC-376 vs Z-FA-FMK, although both could inhibit cathepsin-L with similar efficacy, could be attributed to the higher efficiency of 3CL-Pro inhibition by GC-376 (Fig-2B).

Next, we tested the efficacy of these compounds to inhibit SARS CoV2 delta variant infection in the aforementioned cell lines. Of note, B.1.617.2 delta variant has 30 mutations compared to USA-WA1/2020 isolate of which 9 mutations are in the spike protein region (3). More importantly, one of the mutations is in the polybasic furin cleavage site (P681R) of the spike protein, critical for mediating virus entry. On the other hand, the newly emerged omicron variant has a total of 50 mutations in its genome compared to USA-WA1/2020 and 30 of those mutations exist in the spike gene (4). Therefore, we tested if mutations in newer variants might affect the cathepsin-L cleavage or 3CL-pro cleavage sites and thereby alter the ability of the candidate compounds to inhibit delta/omicron variant infection. As shown in Fig 6A-D, leupeptin was not capable of inhibiting either delta or omicron variant-induced CPE in Vero-TMPRSS2 and Vero-ACE2-TMPRSS2 cells whereas, Z-FA-FMK inhibited CPE in Vero TMPRSS2 cells but not Vero-ACE2-TMPRSS2 cells. Only GC-376 was capable of inhibiting delta variant induced CPE in Vero-ACE2-TMPRSS2 cells similar to the results observed with the USA-WA1/2020 isolate. Noticeably, the efficiency of inhibition of the candidate compounds was consistent between USA-WA1/2020 and the other two variants indicating that the mutations in these variants did not dramatically alter the targeted pathway.

**Figure-6:**
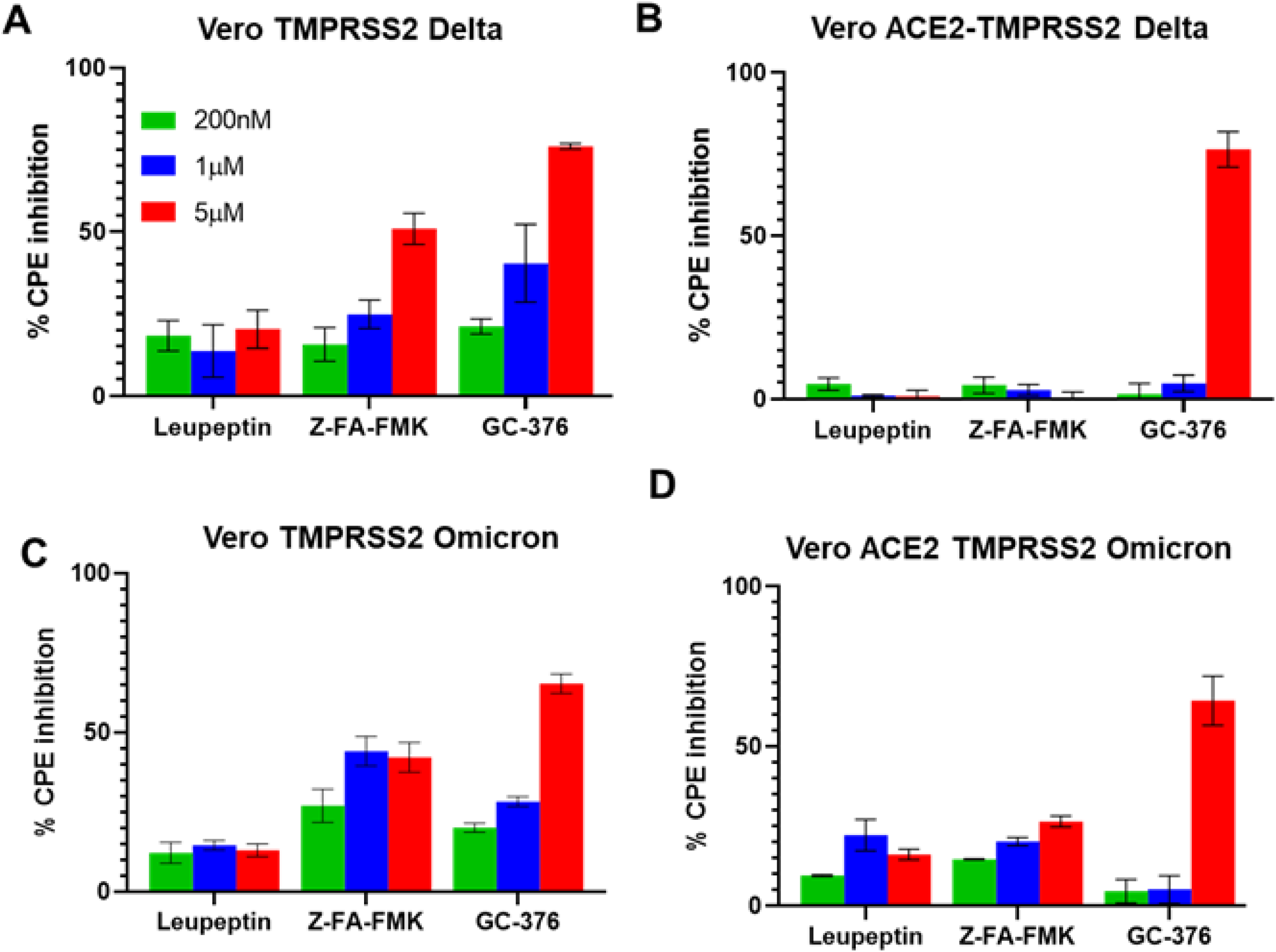
A-B) Effect of post-treatment of leupeptin hemisulfate, Z-FA-FMK and GC-376 on SARS CoV2 B.1.617.2 Delta Variant-induced CPE in (A) Vero-huTMPRSS2 (B) Vero-huACE2-huTMPRSS2 cells against SARS CoV2 B.1.617.2 Delta Variant infection. C-D) Effect of post-treatment of leupeptin hemisulfate, Z-FA-FMK and GC-376 on SARS CoV2 B.1.1.529 omicron-induced CPE in (C) Vero-huTMPRSS2 (D) Vero-huACE2-huTMPRSS2 cells. (Assay was done in triplicates and 2-3 independent experiments. Mean ± SEM values from one independent experiment are shown).

## Discussion

Clinical trial results showed far from optimal activity of remdesivir in inhibiting SARS CoV2 replication (9) and the reasoning for this suboptimal therapeutic efficacy of remdesivir could be that targeting RNA polymerase alone may not be sufficient to clear or control viral load in many severe/hospitalized cases of COVID19. Therefore, targeting additional key steps in SARS CoV2 infection such as virus entry could be critical. Recently, the FDA has granted Emergency Use Authorizations (EUAs) for two drugs namely, molnupiravir and nirmatrelvir/ritonavir for treating COVID19 patients which have shown better efficacy than Remdesivir (10). Two host enzymes, TMPRSS2, and cathepsin-L, are known to be critical for mediating virus entry. Two major virus proteases namely, 3-chymotrypsin-like proteases (3CL-Pro) and papain-like proteases (PL-Pro), are necessary for the processing of several non-structural proteins (NSPs) needed for viral replication, and these virus proteases are attractive targets for antiviral drugs (24, 25). Here, we investigated the key factors regulating SARS CoV2 entry such as cathepsin-L and TMPRSS2 using 293T cells overexpressing human ACE2 (26). Several reports that emerged during the initial phase of the COVID19 pandemic emphasized TMPRSS2 as the most important host factor mediating SARS CoV2 entry and indicated TMPRSS2 inhibition as a therapeutic option for COVID19 (13, 27, 28). Earlier, we had shown that the cathepsin-L cleavage site was conserved in envelope proteins of different families of viruses such as SARS-CoV and Ebola, Hendra, and Nipah viruses, and inhibition of cathepsin-L led to inhibition of entry of these viruses into host cells (18). Using CRISPR gene deletion and pharmacological inhibition, we showed that cathepsin-L could play a more robust role in mediating SARS CoV2 entry into human 293T ACE2 cells than TMPRSS2. These results were consistent with reduced SARS CoV2 induced CPE in *CTSL^Ko^* 293T-ACE2 cells compared to *TMPRSS2™* 293T-ACE2 cells. Similar to our current and past observations, several reports from independent groups have also identified a key role of the cathepsin-L pathway in mediating SARS CoV2 entry (12, 29) (16). Moreover, in recent clinical trials, TMPRSS2 inhibitor camostat mesylate treatment did not affect time to clinical improvement, progression to ICU admission, or mortality (30). Reports are arguing that the use of cathepsin L by coronaviruses such as MERS and 229E is limited mainly to cell lines (31, 32), whereas TMPRSS2 is necessary for the virus spreading and pathogenesis in the infected host (33). In contrast to these observations, a recent study involving COVID19 patients has observed elevated CTSL levels post SARS-CoV-2 infection which directly correlated with disease course and severity (12) suggesting a key role of cathepsin-L in SARS CoV2 infectivity and spread. Similarly, CTSL overexpression was shown to enhance SARS CoV2 pseudovirus entry (12). Hence, unlike other coronaviruses, SARS CoV2 could be more dependent on cathepsin-L than TMPRSS2 for entry into cell.

We screened 5,370 compounds in the TargetMol Bioactives library, a collection of heavily annotated small molecules, including clinically-tested and FDA-approved compounds, and identified 27 compounds with significant cathepsin-L inhibitory activity and potential drug-like properties. Of those 27 compounds, 3 compounds namely, leupeptin hemisulfate, Z-FA-FMK, and GC-376 inhibited SARS CoV2 pseudovirus entry as well as SARS CoV2 induced CPE in 293T ACE2 cells with sub-micromolar IC50 values. More importantly, these drugs inhibited SARS CoV2 induced CPE under both pre-and post-treatment conditions suggesting that the mode of action involved not only a viral entry step but a subsequent step as well. Leupeptin is a reversible inhibitor of trypsin-like proteases and cysteine proteases have been previously shown to prevent SARS CoV1 entry in 293T-ACE2 cells by inhibiting cathepsin-L activity (34). Z-FA-FMK, an irreversible inhibitor of cysteine proteases, has been identified to inhibit SARS CoV2 infection in Vero E6, Caco-2 (human colon adenocarcinoma cells), Huh7 (human hepatocyte carcinoma cells), and LNCaP (human prostate adenocarcinoma cells) and induced Pluripotent Stem Cell-Derived Alveolar Epithelial Cells (iAEC2) as analyzed by morphological cell profiling assay (35). However, this study did not use exogenous ACE2 expression to test the efficacy of the drugs against higher virus titers. GC-376 is a known 3CL-Pro inhibitor, used to treat feline coronavirus infection (36) and has been shown to inhibit SARS CoV2 induced CPE in Vero E6 cells (37). However, the ability of GC-376 to inhibit cathepsin-L and SARS CoV2 entry has not yet been reported. Besides, the efficacy of these molecules against emerging variants of concern such as SARS Cov2 delta and omicron in highly susceptible cell lines has not yet been established.

We found that 293Tcells overexpressing human ACE2 had ~3 logs less susceptibility to SARS CoV2 induced CPE compared to Vero cells overexpressing human TMPRSS2 cells (+ Monkey ACE2). On the other hand, Vero cells overexpressing both human ACE2 and TMPRSS2 were even more susceptible (yielded ~1.5 logs higher virus) susceptible compared to Vero-TMPRSS2 cells. Interestingly, upon subsequent screening of the drug activity in these cell lines, we found that only GC-376 was capable of inhibiting SARS CoV2 induced CPE in all cell lines. Leupeptin which has cathepsin-L inhibitory activity, but lacks 3CL-pro inhibitory activity, inhibited SARS CoV2 CPE only in 293T ACE2 cells and in none of the other cell types. While both Z-FA-FMK and GC-376 had 3CL-Pro inhibitory activity, Z-FA-FMK was capable of inhibiting SARS CoV2 induced CPE in 293T ACE2 and Vero TMPRSS2 cells, but not in Vero ACE2-TMPRSS2 cells. GC-376 which had superior 3CL-Pro activity than Z-FA-FMK was cable of inhibiting SARS CoV2-CPE in all cell lines. These results suggest cathepsin-L inhibition alone may not be sufficient to prevent SARS CoV2 entry in cells that overexpress TMPRSS2. It should be noted that cell type-specific expression of host-proteases with varying levels of expression might be crucial in determining the susceptibility of the host cell types/tissues to SARS CoV2 infection and the efficacy of a given drug. Moreover, differential susceptibility of different cell lines to SARS CoV2 might serve as a surrogate for the differential susceptibility of humans from mild to severe infections, and thus allow for more rigorous validation of drugs in vitro. The importance of choosing an appropriate host cell line can be inferred from the prior knowledge on clinical trials. For instance, initial studies with chloroquine produced promising results in Vero E6 cells (21) but failed in clinical trials (23). However, in subsequent studies, it was proven that chloroquine did not inhibit SARS CoV2 infection in TMPRSS2 overexpressing Vero cells and human lung Calu-3 cells (22) which is consistent with the lack of clinical efficacy of the drug. In another instance, the failure of TMPRSS inhibitor in clinical trials underscores the insufficiency of targeting a single mode of action, which can be readily overcome by fulminant infection in hospitalized patients (30).

Independent studies have shown attenuated neutralizing activity of the sera from convalescent patients and individuals vaccinated with either Pfizer or Moderna or AstraZeneca vaccine against delta and omicron variants compared to the prototypic (USA-WA1/2020) strain (3, 38) (4, 5). It should be noted that all these vaccines target the spike protein region of the Wuhan strain. Delta variant mutations in the spike region involve the receptor-binding domain (RBD), the N-terminal antigenic supersite, and the polybasic furin cleavage site (P681R) (3). The newer mutations in the spike protein regions could contribute to immune evasion by viral variants including the delta and omicron variants. The efficacy of leupeptin hemisulfate, Z-FA-FMK, and GC-376 against delta variant in different cell lines used in this study was comparable to their effects on the USA- WA1/2020 isolate and thus suggested that the observed mutations didn’t affect the drug targets. Thus, we report GC-376 can be a potential drug candidate for treating SARS CoV2 including the delta and omicron variant. Given that the 3CL-Pro is targeted by the nirmatrelvir(39), it is possible that this drug can be effective against the omicron variant as well but, yet to be proven.

GC-376 (a prodrug of GC-373) is structurally similar to nirmatrelvir (PF-07321332). Nirmatrelvir is an analog with a substitution of aldehyde covalent cysteine (acceptor) with a nitrile. This drug is co-administered with low dose ritonavir (used for blocking CYP3A4 to slow metabolism of nirmatrelvir) (39). The GC-376 and nimratrelvir and their prodrugs are SARS-CoV-2 virus 3CL protease inhibitors. Core structures overlay very well and are essential for 3CL-pro inhibition. Given that the 3CL-Pro is targeted by the nirmatrelvir, it is possible that this drug can be effective against the omicron variant as well but, yet to be proven. Interestingly a recent study, using the K18-hACE2 mouse model, has shown antiviral efficacy of GC-376 against SARS-CoV-2(40). GC-376 treatment was not toxic and improved survival from 0 to 20% in mice challenged with this virus. Treatment also led to reduced viral loads and reduced inflammation in comparison to controls. This study with SARS-CoV2 further supports the notion that GC-376 is a promising lead candidate for further development.

Here, we report GC-376 can be a potential drug candidate for treating SARS CoV2 including the delta and omicron variants. We furthermore report that GC-376 can inhibit SARS CoV2 infection by inhibiting more than one proteolytic enzyme, thus limiting the ability of genetic variants to escape their anti-viral effect. Taken together, our findings highlight the importance of targeting more than one aspect of the virus life cycle and choosing a broad range of host cell lines for anti-viral drug screening.

## Acknowledgments

This work was supported by funds from Vanda Pharmaceuticals Inc, USA to Dr. BP.

## References

1. Li Q, Guan X, Wu P, Wang X, Zhou L, Tong Y, Ren R, Leung KSM, Lau EHY, Wong JY, Xing X, Xiang N, Wu Y, Li C, Chen Q, Li D, Liu T, Zhao J, Liu M, Tu W, Chen C, Jin L, Yang R, Wang Q, Zhou S, Wang R, Liu H, Luo Y, Liu Y, Shao G, Li H, Tao Z, Yang Y, Deng Z, Liu B, Ma Z, Zhang Y, Shi G, Lam TTY, Wu JT, Gao GF, Cowling BJ, Yang B, Leung GM, Feng Z. 2020. Early Transmission Dynamics in Wuhan, China, of Novel Coronavirus-Infected Pneumonia. N Engl J Med 382:1199–1207.

2. Lopez Bernal J, Andrews N, Gower C, Gallagher E, Simmons R, Thelwall S, Stowe J, Tessier E, Groves N, Dabrera G, Myers R, Campbell CNJ, Amirthalingam G, Edmunds M, Zambon M, Brown KE, Hopkins S, Chand M, Ramsay M. 2021. Effectiveness of Covid-19 Vaccines against the B.1.617.2 (Delta) Variant. N Engl J Med doi:10.1056/NEJMoa2108891.

3. Edara VV, Pinsky BA, Suthar MS, Lai LL, Davis-Gardner ME, Floyd K, Flowers MW, Wrammert J, Hussaini L, Ciric CR, Bechnak S, Stephens K, Graham BS, Mokhtari EB, Mudvari P, Boritz E, Creanga A, Pegu A, Derrien-Colemyn A, Henry AR, Gagne M, Douek DC, Sahoo MK, Sibai M, Solis D, Webby RJ, Jeevan T, Fabrizio TP. 2021. Infection and Vaccine-Induced Neutralizing-Antibody Responses to the SARS-CoV-2 B.1.617 Variants. New England Journal of Medicine 385:664–666.

4. Dejnirattisai W, Huo J, Zhou D, Zahradnik J, Supasa P, Liu C, Duyvesteyn HME, Ginn HM, Mentzer AJ, Tuekprakhon A, Nutalai R, Wang B, Dijokaite A, Khan S, Avinoam O, Bahar M, Skelly D, Adele S, Johnson SA, Amini A, Ritter T, Mason C, Dold C, Pan D, Assadi S, Bellass A, Omo-Dare N, Koeckerling D, Flaxman A, Jenkin D, Aley PK, Voysey M, Costa Clemens SA, Naveca FG, Nascimento V, Nascimento F, Fernandes da Costa C, Resende PC, Pauvolid-Correa A, Siqueira MM, Baillie V, Serafin N, Ditse Z, Silva KD, Madhi S, Nunes MC, Malik T, Openshaw PJ, Baillie JK, Semple MG, et al. 2021. Omicron-B.1.1.529 leads to widespread escape from neutralizing antibody responses. bioRxiv doi:10.1101/2021.12.03.471045.

5. Syed AM, Ciling A, Khalid MM, Sreekumar B, Chen PY, Kumar GR, Silva I, Milbes B, Kojima N, Hess V, Shacreaw M, Lopez L, Brobeck M, Turner F, Spraggon L, Taha TY, Tabata T, Chen IP, Ott M, Doudna JA. 2022. Omicron mutations enhance infectivity and reduce antibody neutralization of SARS-CoV-2 virus-like particles. medRxiv doi:10.1101/2021.12.20.21268048.

6. Takashita E, Kinoshita N, Yamayoshi S, Sakai-Tagawa Y, Fujisaki S, Ito M, Iwatsuki-Horimoto K, Chiba S, Halfmann P, Nagai H, Saito M, Adachi E, Sullivan D, Pekosz A, Watanabe S, Maeda K, Imai M, Yotsuyanagi H, Mitsuya H, Ohmagari N, Takeda M, Hasegawa H, Kawaoka Y. 2022. Efficacy of Antibodies and Antiviral Drugs against Covid-19 Omicron Variant. N Engl J Med doi:10.1056/NEJMc2119407.

7. Rajpoot S, Alagumuthu M, Baig MS. 2021. Dual targeting of 3CL(pro) and PL(pro) of SARS-CoV-2: A novel structure-based design approach to treat COVID-19. Curr Res Struct Biol 3:9–18.

8. Hillen HS, Kokic G, Farnung L, Dienemann C, Tegunov D, Cramer P. 2020. Structure of replicating SARS-CoV-2 polymerase. Nature 584:154–156.

9. Beigel JH, Tomashek KM, Dodd LE, Mehta AK, Zingman BS, Kalil AC, Hohmann E, Chu HY, Luetkemeyer A, Kline S, de Castilla DL, Finberg RW, Dierberg K, Tapson V, Hsieh L, Patterson TF, Paredes R, Sweeney DA, Short WR, Touloumi G, Lye DC, Ohmagari N, Oh MD, Ruiz-Palacios GM, Benfield T, Fatkenheuer G, Kortepeter MG, Atmar RL, Creech CB, Lundgren J, Babiker AG, Pett S, Neaton JD, Burgess TH, Bonnett T, Green M, Makowski M, Osinusi A, Nayak S, Lane HC, Grp A-S. 2020. Remdesivir for the Treatment of Covid-19-Final Report. New England Journal of Medicine 383:1813–1826.

10. Cully M. 2022. A tale of two antiviral targets - and the COVID-19 drugs that bind them. Nature Reviews Drug Discovery 21:3–5.

11. Ren SY, Wang WB, Gao RD, Zhou AM. 2022. Omicron variant (B.1.1.529) of SARS-CoV-2: Mutation, infectivity, transmission, and vaccine resistance. World J Clin Cases 10:1–11.

12. Zhao MM, Yang WL, Yang FY, Zhang L, Huang WJ, Hou W, Fan CF, Jin RH, Feng YM, Wang YC, Yang JK. 2021. Cathepsin L plays a key role in SARS-CoV-2 infection in humans and humanized mice and is a promising target for new drug development. Signal Transduction and Targeted Therapy 6.

13. Hoffmann M, Kleine-Weber H, Schroeder S, Kruger N, Herrler T, Erichsen S, Schiergens TS, Herrler G, Wu NH, Nitsche A, Muller MA, Drosten C, Pohlmann S. 2020. SARS-CoV-2 Cell Entry Depends on ACE2 and TMPRSS2 and Is Blocked by a Clinically Proven Protease Inhibitor. Cell 181:271–+.

14. Heald-Sargent T, Gallagher T. 2012. Ready, Set, Fuse! The Coronavirus Spike Protein and Acquisition of Fusion Competence. Viruses-Basel 4:557–580.

15. Zhu YK, Feng F, Hu GW, Wang YY, Yu Y, Zhu YF, Xu W, Cai X, Sun ZP, Han WD, Ye R, Qu D, Ding Q, Huang XX, Chen HJ, Xu W, Xie YH, Cai QL, Yuan ZH, Zhang R. 2021. A genome-wide CRISPR screen identifies host factors that regulate SARS-CoV-2 entry. Nature Communications 12.

16. Jaimes JA, Millet JK, Whittaker GR. 2020. Proteolytic Cleavage of the SARS-CoV-2 Spike Protein and the Role of the Novel S1/S2 Site. Iscience 23.

17. Millet JK, Whittaker GR. 2014. Host cell entry of Middle East respiratory syndrome coronavirus after two-step, furin-mediated activation of the spike protein. Proceedings of the National Academy of Sciences of the United States of America 111:15214–15219.

18. Elshabrawy HA, Fan JL, Haddad CS, Ratia K, Broder CC, Caffrey M, Prabhakar BS. 2014. Identification of a Broad-Spectrum Antiviral Small Molecule against Severe Acute Respiratory Syndrome Coronavirus and Ebola, Hendra, and Nipah Viruses by Using a Novel High-Throughput Screening Assay. Journal of Virology 88:4353–4365.

19. Dahlin JL, Nissink JW, Strasser JM, Francis S, Higgins L, Zhou H, Zhang Z, Walters MA. 2015. PAINS in the assay: chemical mechanisms of assay interference and promiscuous enzymatic inhibition observed during a sulfhydryl-scavenging HTS. J Med Chem 58:2091–2113.

20. Xia Z, Sacco M, Hu Y, Ma C, Meng X, Zhang F, Szeto T, Xiang Y, Chen Y, Wang J. 2021. Rational Design of Hybrid SARS-CoV-2 Main Protease Inhibitors Guided by the Superimposed Cocrystal Structures with the Peptidomimetic Inhibitors GC-376, Telaprevir, and Boceprevir. ACS Pharmacol Transl Sci 4:1408–1421.

21. Liu J, Cao RY, Xu MY, Wang X, Zhang HY, Hu HR, Li YF, Hu ZH, Zhong W, Wang ML. 2020. Hydroxychloroquine, a less toxic derivative of chloroquine, is effective in inhibiting SARS-CoV-2 infection in vitro. Cell Discovery 6.

22. Hoffmann M, Mosbauer K, Hofmann-Winkler H, Kaul A, Kleine-Weber H, Kruger N, Gassen NC, Muller MA, Drosten C, Pohlmann S. 2020. Chloroquine does not inhibit infection of human lung cells with SARS-CoV-2. Nature 585:588–590.

23. Ho TC, Wang YH, Chen YL, Tsai WC, Lee CH, Chuang KP, Chen YA, Yuan CH, Ho SY, Yang MH, Tyan YC. 2021. Chloroquine and Hydroxychloroquine: Efficacy in the Treatment of the COVID-19. Pathogens 10.

24. Mody V, Ho J, Wills S, Mawri A, Lawson L, Ebert MCCJC, Fortin GM, Rayalam S, Taval S. 2021. Identification of 3-chymotrypsin like protease (3CLPro) inhibitors as potential anti-SARS-CoV-2 agents. Communications Biology 4.

25. Shin D, Mukherjee R, Grewe D, Bojkova D, Baek K, Bhattacharya A, Schulz L, Widera M, Mehdipour AR, Tascher G, Geurink PP, Wilhelm A, van Noort GJV, Ovaa H, Muller S, Knobeloch KP, Rajalingam K, Schulman BA, Cinatl J, Hummer G, Ciesek S, Dikic I. 2020. Papain-like protease regulates SARS-CoV-2 viral spread and innate immunity. Nature 587:657–+.

26. Padmanabhan P, Desikan R, Dixit NM. 2020. Targeting TMPRSS2 and Cathepsin B/L together may be synergistic against SARS-CoV-2 infection. Plos Computational Biology 16.

27. Hu X, Shrimp JH, Guo H, Xu M, Chen CZ, Zhu W, Zakharov AV, Jain S, Shinn P, Simeonov A, Hall MD, Shen M. 2021. Discovery of TMPRSS2 Inhibitors from Virtual Screening as a Potential Treatment of COVID-19. Acs Pharmacology & Translational Science 4:1124–1135.

28. McKee DL, Sternberg A, Stange U, Laufer S, Naujokat C. 2020. Candidate drugs against SARS-CoV-2 and COVID-19. Pharmacological Research 157.

29. Ou X, Liu Y, Lei X, Li P, Mi D, Ren L, Guo L, Guo R, Chen T, Hu J, Xiang Z, Mu Z, Chen X, Chen J, Hu K, Jin Q, Wang J, Qian Z. 2020. Characterization of spike glycoprotein of SARS-CoV-2 on virus entry and its immune crossreactivity with SARS-CoV. Nat Commun 11:1620.

30. Gunst JD, Staerke NB, Pahus MH, Kristensen LH, Bodilsen J, Lohse N, Dalgaard LS, Bronnum D, Frobert O, Honge B, Johansen IS, Monrad I, Erikstrup C, Rosendal R, Vilstrup E, Mariager T, Bove DG, Offersen R, Shakar S, Cajander S, Jorgensen NP, Sritharan SS, Breining P, Jespersen S, Mortensen KL, Jensen ML, Kolte L, Frattari GS, Larsen CS, Storgaard M, Nielsen LP, Tolstrup M, Saedder EA, Ostergaard LJ, Ngo HTT, Jensen MH, Hojen JF, Kjolby M, Sogaard OS. 2021. Efficacy of the TMPRSS2 inhibitor camostat mesilate in patients hospitalized with Covid-19-a double-blind randomized controlled trial. Eclinicalmedicine 35.

31. Shirato K, Kawase M, Matsuyama S. 2018. Wild-type human coronaviruses prefer cell-surface TMPRSS2 to endosomal cathepsins for cell entry. Virology 517:9–15.

32. Shirato K, Kanou K, Kawase M, Matsuyama S. 2017. Clinical Isolates of Human Coronavirus 229E Bypass the Endosome for Cell Entry. Journal of Virology 91.

33. Iwata-Yoshikawa N, Okamura T, Shimizu Y, Hasegawa H, Takeda M, Nagata N. 2019. TMPRSS2 Contributes to Virus Spread and Immunopathology in the Airways of Murine Models after Coronavirus Infection. Journal of Virology 93.

34. Simmons G, Gosalia DN, Rennekamp AJ, Reeves JD, Diamond SL, Bates P. 2005. Inhibitors of cathepsin L prevent severe acute respiratory syndrome coronavirus entry. Proc Natl Acad Sci U S A 102:11876–11881.

35. Mirabelli C, Wotring JW, Zhang CJ, McCarty SM, Fursmidt R, Pretto CD, Qiao Y, Zhang Y, Frum T, Kadambi NS, Amin AT, O’Meara TR, Spence JR, Huang J, Alysandratos KD, Kotton DN, Handelman SK, Wobus CE, Weatherwax KJ, Mashour GA, O’Meara MJ, Chinnaiyan AM, Sexton JZ. 2021. Morphological cell profiling of SARS-CoV-2 infection identifies drug repurposing candidates for COVID-19. Proc Natl Acad Sci U S A 118.

36. Pedersen NC, Kim Y, Liu HW, Kankanamalage ACG, Eckstrand C, Groutas WC, Bannasch M, Meadows JM, Chang KO. 2018. Efficacy of a 3C-like protease inhibitor in treating various forms of acquired feline infectious peritonitis. Journal of Feline Medicine and Surgery 20:378–392.

37. Hu YM, Ma CL, Szeto T, Hurst B, Tarbet B, Wang J. 2021. Boceprevir, Calpain Inhibitors II and XII, and GC-376 Have Broad-Spectrum Antiviral Activity against Coronaviruses. Acs Infectious Diseases 7:586–597.

38. Planas D, Veyer D, Baidaliuk A, Staropoli I, Guivel-Benhassine F, Rajah MM, Planchais C, Porrot F, Robillard N, Puech J, Prot M, Gallais F, Gantner P, Velay A, Le Guen J, Kassis-Chikhani N, Edriss D, Belec L, Seve A, Courtellemont L, Pere H, Hocqueloux L, Fafi-Kremer S, Prazuck T, Mouquet H, Bruel T, Simon-Loriere E, Rey FA, Schwartz O. 2021. Reduced sensitivity of SARS-CoV-2 variant Delta to antibody neutralization. Nature 596:276–280.

39. Owen DR, Allerton CMN, Anderson AS, Aschenbrenner L, Avery M, Berritt S, Boras B, Cardin RD, Carlo A, Coffman KJ, Dantonio A, Di L, Eng H, Ferre R, Gajiwala KS, Gibson SA, Greasley SE, Hurst BL, Kadar EP, Kalgutkar AS, Lee JC, Lee J, Liu W, Mason SW, Noell S, Novak JJ, Obach RS, Ogilvie K, Patel NC, Pettersson M, Rai DK, Reese MR, Sammons MF, Sathish JG, Singh RSP, Steppan CM, Stewart AE, Tuttle JB, Updyke L, Verhoest PR, Wei L, Yang Q, Zhu Y. 2021. An oral SARS-CoV-2 M(pro) inhibitor clinical candidate for the treatment of COVID-19. Science 374:1586–1593.

40. Caceres CJ, Cardenas-Garcia S, Carnaccini S, Seibert B, Rajao DS, Wang J, Perez DR. 2021. Efficacy of GC-376 against SARS-CoV-2 virus infection in the K18 hACE2 transgenic mouse model. Sci Rep 11:9609.

